# Exploring Mechanisms of Glucose Uptake Regulation and Dilution Resistance in Growing Cancer Cells

**DOI:** 10.1101/2020.01.02.892729

**Authors:** Daniel M. Tveit, Gunhild Fjeld, Tormod Drengstig, Fabian V. Filipp, Peter Ruoff, Kristian Thorsen

**Affiliations:** Department of Electrical Engineering and Computer Science, University of Stavanger, Stavanger, Norway; Centre for Organelle Research, University of Stavanger, Stavanger, Norway; Systems Biology and Cancer Metabolism, Program for Quantitative Systems Biology, University of California Merced, Merced, CA, USA

## Abstract

Most cancer cells rely on aerobic glycolysis and increased glucose uptake for the production of biosynthetic precursors needed to support rapid proliferation. Increased glucose uptake and glycolytic activity may result in intracellular acidosis and increase of osmotically active substances, leading to cell swelling. This causes dilution of cellular constituents, which can markedly influence cellular reactions and the function of proteins, and hence, control mechanisms used by cancer cells to maintain a highly glycolytic phenotype must be robust to dilution. In this paper, we review the literature on cancer cell metabolism and glucose uptake, and employ mathematical modeling to examine control mechanisms in cancer cell metabolism that show robust homeostatic control in the presence of dilution. Using differential gene expression data from the Expression Atlas database, we identify the key components of glucose uptake in cancer, in order to guide the construction of a mathematical model. By simulations of this model we show that while negative feedback from downstream glycolytic metabolites to glucose transporters is sufficient for homeostatic control of glycolysis in a constant cellular volume, it is necessary to control intermediate glycolytic enzymes in order to achieve homeostatic control during growth. With a focus on glucose uptake in cancer, we demonstrate a systems biology approach to the identification, reduction, and analysis of complex regulatory systems.

**SIGNIFICANCE:** Rapid proliferation and increased glycolytic activity in cancer cells lead to dilution of cellular constituents, which can markedly influence cellular reactions and the function of proteins. Therefore, control mechanisms used by cancer cells to maintain a highly glycolytic phenotype must be robust to dilution. We construct a mathematical model of glucose uptake in cancer, and using a systems biology approach to the analysis of regulatory networks, identify the presence of integral control motifs as a means for achieving dilution resistance. Furthermore, we show that while negative feedback from downstream glycolytic metabolites to glucose transporters is sufficient for homeostatic control of glycolysis in a constant cellular volume, it is necessary to control intermediate glycolytic enzymes to achieve homeostatic control during growth.

## INTRODUCTION

It is well established that cell swelling and shrinkage affect important cellular functions, in part by dilution and concentration of cellular compounds (1–4). Such changes in concentration can markedly influence the function of intracellular proteins (1). This has been demonstrated in studies on the effect of volume change on enzyme reactions in solitary vesicles, showing that there is a significant impact on the dynamical and steady state behavior of these reactions (5). The significance of dilution due to growth is emphasized by the cell-size control mechanism employed in budding-yeast. In these cells, the concentration of a cell cycle activator maintained at a constant level during growth relative to a growth diluted inhibitor provides a measurement of cell volume and a molecular mechanism for cell-size control (6, 7). Although most proteins are maintained at constant concentrations in growing cells, owing to mRNA amounts and number of ribosomes increasing with cell size, we lack an understanding of the molecular mechanisms that coordinate biosynthesis to achieve constant protein concentrations during growth (7, 8). Investigations into the performance of so-called integral control motifs (ICMs) have revealed mechanisms by which robustness to dilution can be achieved (9, 10). In this paper, we look into homeostatic mechanisms regulating glucose uptake in rapidly growing cancer cells. We identify the presence of ICMs as part of glucose uptake in cancer, and investigate how homeostatic control of metabolite and protein concentrations is achieved in the presence of dilution due to growth.

Most cancer cells show an increased uptake and metabolism of glucose, a phenotype that can be detected by ^18^fluoro-deoxyglucose positron emission tomography (FDG-PET) (11–13). This growth mode relies on a balanced production of cellular components to avoid molecular crowding and solvent capacity constraints (14, 15). The cell represents a tiny reagent reservoir and is reliant on a balanced influx and efflux of compounds to support growth rates corresponding to that seen in cancer. Thus, as the cell expands, its constituents need to increase at the same rate to meet the growth requirements, meaning a proportional increase in nucleic acids, polysaccharides, proteins, and lipids (16). Aside from biomass formation for the purpose of growth, metabolism also affects cell volume through uptake of nutrients, by creation of osmotically active substances, developing intracellular acidosis, and by depletion of available ATP (1, 17, 18). In fact, increased cellular volume appears to be required for proliferation, and hypertonic shrinkage inhibits cell proliferation, whereas slight osmotic swelling has the opposite effect (1). In contrast, differentiation is followed by cell shrinkage in a number of cells (1).

In the following, we take a look at some key elements of the rewiring of glycolysis that produce the increased glycolytic activity seen in cancer. Next, we look at the various control mechanisms in place that maintain this increased glucose uptake and metabolic activity. We then focus our attention to glucose uptake, and show that differential gene expression of cancer and normal cells corroborate the reported rewiring in cancer. With this information, we construct a mathematical model of glucose uptake in cancer, formulated as a system of ordinary differential equations (ODEs). We construct the model in a stepwise manner, where each step adds a layer of regulation to the model. This is done in order to investigate the role each control mechanism serve. We show how dilution is incorporated into the model, and run simulations for each step of model construction. Finally, we show how control mechanisms of glucose uptake in cancer form ICMs, and how this enables robust homeostatic control of glycolysis, even in the presence of dilution.

### Rewiring of Glycolysis in Cancer

Cancer cells show an increased reliance on glycolysis and lactic acid fermentation, even in the presence of oxygen, and a more glycolytic phenotype is persistent with a more aggressive cancer cell type (12, 19, 20). This is known as the Warburg effect, or aerobic glycolysis, and is necessary in order to meet the increased demands of rapid proliferation (11). In cancer cells, the Warburg effect is in supplement to oxidative phosphorylation rather than a replacement (21). This is in contrast to normal cells that maintain a high rate of glycolysis at the expense of oxidative phosphorylation; a phenomenon known as the Crabtree effect (21). However, in the hypoxic tumor microenvironment, cancer cells naturally show a decreased reliance on oxidative phosphorylation (21, 22). The increased glycolytic flux in cancer supplies biosynthetic pathways with precursors, meets the increased bioenergetic demand of proliferation, and contributes to tumor invasion through the excretion of lactate and consequent acidification of the tumor microenvironment (11, 12, 21, 23, 24). The mechanisms that reprogram metabolism in cancer are often cancer-specific, nevertheless, there are common hallmarks, notably a shift towards protein isoforms that promote biosynthesis and proliferation (11, 21).

In the first step of glycolysis, glucose is transported into the cell. The GLUT (gene symbol *SLC2A*) family of glucose transporters are membrane-spanning proteins facilitating the transport of sugars across biological membranes along the concentration gradient (25, 26). GLUT1 is one of 14 currently identified GLUT proteins expressed in humans, and is expressed in almost every tissue (27–30). Together with its high affinity for glucose, this gives GLUT1 a clear role in the basal glucose uptake of most tissues (25, 28, 29). Elevated expression of GLUT1 has been reported in most cancers, and the expression level correlates reciprocally with the survival of cancer patients (12, 23, 30). Hypoxia-inducible factor-1 (HIF-1), a dimer of HIF-1α and HIF-1β, is one of the factors responsible for upregulating GLUT1 in tumor cells (12, 21, 30, 31). HIF-1β is constitutively expressed, whereas HIF-1α is regulated through oxygen-dependent and oxygen-independent mechanisms (31). GLUT1 expression is upregulated through hypoxia-response elements on the GLUT1 promoter that bind HIF-1 (30). HIF-1α has increased levels in most cancers, which provides a mechanism by which cancer cells overexpress GLUT1 (12, 30, 31). Other factors known to cause overexpression and translocation of GLUT1 to the cell membrane in cancer include the oncoprotein c-Myc, protein kinase Akt/PKB, and oncogenic KRAS and BRAF (12, 21, 30).

Glycolysis consists of several reversible reactions and three (essentially) irreversible reactions (see Figure 1). Because they are irreversible, these three reactions represent committed steps of glycolysis, and the enzymes that drive these reactions function as gatekeepers of glycolysis and have a key role in regulating the glycolytic flux (21). In the first irreversible reaction of glycolysis, glucose is phosphorylated to glucose 6-phosphate (G6P) by hexokinase, coupled to the dephosphorylation of ATP (13, 32, 33). Hexokinase 2 (HK2) is one of four isoforms of hexokinase found in mammalian tissue (13). HK2 has a very high affinity for glucose, with a Michaelis constant (*K*_M_ value) of 0.02–0.03 mm (13, 32). To support increased glucose uptake in cancer, HK2 is overexpressed and bound to the outer mitochondrial membrane protein voltage-dependent anion channel (VDAC) (13, 21, 32). VDAC supplies HK2 with ATP by recruiting help from ATP synthase and adenine nucleotide translocator, resulting in a mechanism that rapidly converts glucose to G6P (13). HK2 is product inhibited by G6P, however, it is likely that this inhibition is minimal due to rapid utilization of G6P in cancer cells (21, 32).

**Figure 1:**
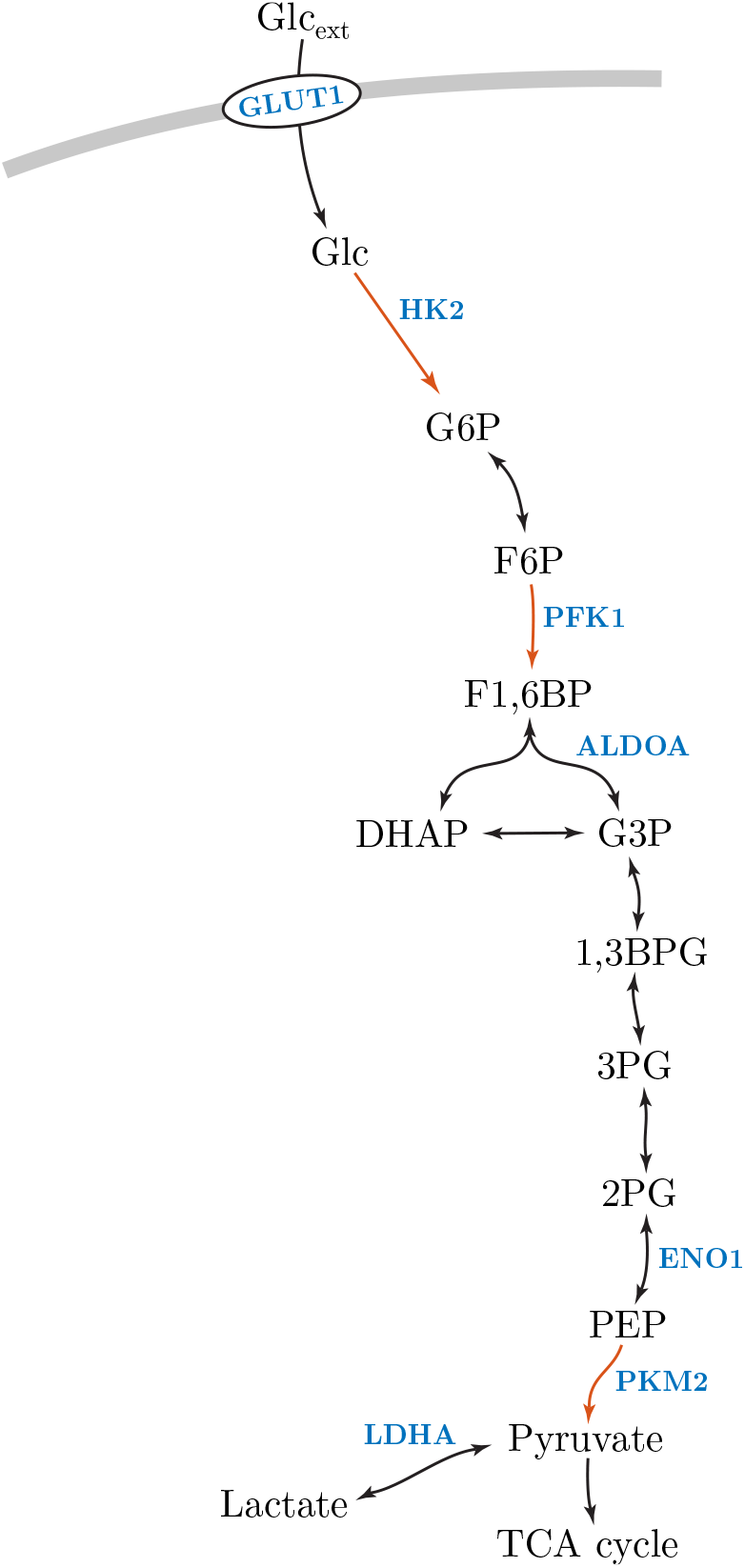
Schematic of glycolysis with some of the commonly overexpressed isoforms in cancer highlighted in blue. Reactions highlighted in red indicate the committed steps of glycolysis. Abbreviations: Extracellular glucose (Glc_ext_), glucose transporter 1 (GLUT1), glucose (Glc), hexokinase 2 (HK2), glucose 6-phosphate (G6P), fructose 6-phosphate (F6P), phosphofructokinase 1 (PFK1), fructose 1,6-bisphosphate (F1,6BP), aldolase A (ALDOA), dihydroxyacetone phosphate (DHAP), glyceraldehyde 3- phosphate (G3P), 1,3-bisphosphoglycerate (1,3BPG), 3-phosphoglycerate (3PG), 2-phosphoglycerate (2PG), enolase 1 (ENO1), phosphoenolpyruvate (PEP), pyruvate kinase M2 (PKM2), lactate dehydrogenase A (LDHA), tricarboxylic acid cycle (TCA cycle).

The second irreversible reaction of glycolysis is catalyzed by phosphorfructokinase 1 (PFK1), and is the phosphorylation of fructose 6-phosphate (F6P) to fructose 1,6-bisphosphate (F1,6BP) with the concomitant dephosphorylation of ATP (21, 33, 34). PFK1 is a tetrameric enzyme that exists in liver (PFKL), muscle (PFKM), and platelet (PFKP) isoforms in mammalian cells (21, 34, 35). PFK1 expression is upregulated in cancer cells, and increased expression of the PFKP isoform is a characteristic feature of cancer (34, 35). Krüppel-like factor 4 (KLF4), which has elevated levels in certain cancer types, has been shown to activate transcription of the *PFKP* gene by directly binding to its promoter (34). In addition, PFK1 is allosterically activated by fructose 2,6-bisphosphate (F2,6BP), which shows increased generation associated with overexpression of the phosphofructokinase 2 (PFK2) isoform PFKFB3 in cancer (21).

The third irreversible reaction of glycolysis is the conversion of phosphoenolpyruvate (PEP) to pyruvate by the transfer of a phosphoryl group to ADP (21, 33). Cancer cells control this reaction by expressing the low-affinity M2 isoform of pyruvate kinase (PKM2) (11, 21). The PKM2 tetramer is allosterically regulated by various metabolites and responds to nutritional and stress signals, whereas the normal M1 isoform of pyruvate kinase (PKM1) is a constitutively active tetramer (21, 36). The regulation of PKM2 enables cancer cells to dictate the flow of carbon into biosynthetic pathways and adapt to different conditions of nutrient availability and anabolic demands (11, 21, 36). Additionally, PKM2 is regulated between its metabolically active tetrameric form and metabolically inactive dimeric form, where the PKM2 dimer is imported into the nucleus and stimulates transcription of glycolytic genes (36).

In addition to the key regulatory enzymes of glycolysis described above, other important glycolytic enzymes are also upregulated in cancer. For example, of the lactate dehydrogenases (LDHs), LDHA is the predominantly expressed isozyme in cancer (21). LDHA has a high affinity for pyruvate, and favors the conversion of pyruvate to lactate (21). Enolase 1 (ENO1) is induced in cancer cells through HIF-1α overexpression (30, 31). Aldolase A (ALDOA) is the predominant aldolase isoform expressed in hepatoma and gastric cancer tissues, and favors the cleaving of F1,6BP (21, 37). Taken together, the glycolytic isoforms expressed in cancer show a concerted effort to increase glycolytic activity and promote production of biosynthetic precursors. A schematic of glycolysis is shown in Figure 1, highlighting some of the key isoforms that are commonly overexpressed in cancer.

### Regulation of Glucose Uptake in Cancer

We now focus our attention to glucose uptake and the initial steps of glycolysis, and discuss the control mechanisms that regulate glucose uptake in cancer. Although key glycolytic enzymes are upregulated in cancer, they are still involved in metabolic regulation and respond to signals such as nutritional and oxidative stress, however, this regulation changes to favor proliferation (11, 21, 36, 38). Regulation of nutrient transporters by the availability of nutrients is a phenomenon observed in bacteria and yeast, and similarly, an inhibitory effect of glucose on GLUT1 expression has been observed in several mammalian cell lines (39, 40). To study the effect of glucose on GLUT1 expression, cells have been subjected to glucose deprivation experiments, with the common result that GLUT1 content at the cell surface is increased (39, 41–45). This is achieved by different mechanisms, including increased GLUT1 mRNA transcription and stability, increased protein synthesis or decreased protein degradation, and translocation of the transporter to the cell membrane (39).

Extracellular glucose supply directly affects the intracellular glucose level (46). Thus, it is possible that GLUT1 content at the cell surface is regulated in some way by the intracellular level of glucose, as has been previously suggested (41, 47). In fact, comparisons of mammary tumors and normal mammary tissue in mice have shown that increased GLUT1 level correlates with decreased intracellular glucose level and increased glycolytic activity (38). One way in which intracellular glucose affects GLUT1 expression is via AMP-activated protein kinase (AMPK) (48). AMPK is comprised of one catalytic α-subunit, and two regulatory subunits, β and γ (48, 49). Intracellular glucose regulates AMPK activity in a few different ways: An abundance of glucose will quickly be phosphorylated to G6P by HK2. G6P is then used to supply glycolysis, lowering the AMP/ATP and ADP/ATP ratios, keeping AMPK from being activated by the binding of AMP and ADP (48). High glucose levels and increased biomass generation also reduce the NAD^+^/NADH ratio, which indirectly inhibits AMPK through silent information regulator T1 (SIRT1) and serine-threonine liver kinase B1 (LKB1) (21, 48, 50, 51). Downstream of G6P, the accumulation of diacylglycerol (DAG) and glycogen both lead to inhibition of AMPK. DAG inhibits AMPK by activating protein kinase C (PKC), which in turn induces the inhibitory phosphorylation of the AMPK α-subunit, while glycogen inhibits AMPK by binding to the β-subunit (48). In addition, activation of protein phosphatase 2A (PP2A) as a result of high glucose levels inhibits AMPK (48, 52, 53).

AMPK in turn has been shown to affect GLUT1 expression (54). One mechanism by which this happens is by increasing the degradation of thioredoxin interacting protein (TXNIP). TXNIP can bind directly to GLUT1 and induce internalization, as well as reduce GLUT1 mRNA level (48, 55). Another suggested mechanism is that downstream of AMPK, p38 mitogen-activated protein kinase (MAPK) activation leads to enhancement of GLUT1 mediated glucose transport (56).

Another important aspect of glucose uptake is the regulation of HK2, as it drives the first committed step of glycolysis and maintains a high concentration gradient of glucose across the cell membrane, thereby driving the facilitated diffusion of glucose by GLUT1 (21, 57, 58). Activators of the HK2 promoter include glucose, insulin, glucagon, p53, cAMP, and hypoxic conditions (13, 32, 59). Interestingly, it is glucose rather than downstream glycolytic metabolites that activate the HK2 promoter (13, 32, 59–61). Together with the fact that HK2 phosphorylates glucose to G6P in a reaction that is essentially irreversible, these two compounds form a stabilizing feedback connection (13, 62). Additionally, the binding of HK2 to the outer mitochondrial membrane via VDAC helps prevent apoptosis in cancer cells (13, 32). Together with the diminished inhibition (or possibly saturated inhibition) of HK2 by G6P that is associated with mitochondrial bound HK2, this gives a clear role for HK2 in promoting a malignant phenotype (32, 63–65).

The control mechanisms discussed above are summarized in Figure 2A. Here, glucose uptake and supply to metabolism includes regulatory pathways that inhibit GLUT1 mediated glucose uptake via AMPK, as well as the stabilizing feedback connection formed by glucose and HK2. The mechanisms that affect AMPK depend on the production of G6P, and therefore, G6P represents the potential for these mechanisms to ultimately affect GLUT1 mediated glucose uptake. Before we can construct a mathematical model of the system in Figure 2A, activating and inhibiting pathways need to be translated into reactions that can be described by reaction kinetic equations. To this end, parallel pathways with similar overall effects are grouped together, shown in Figure 2B. These combined pathways are then turned into activating or inhibiting reactions affecting generation or removal reactions of the compounds considered, shown in Figure 2C. The conversion of the system in Figure 2B to the system in Figure 2C preserves the effect one compound has on another, however, this conversion is not unique. For example, a negative effect of G6P on GLUT1 content at the cell surface could also be achieved if G6P activates the degradation or internalization of GLUT1 (62). Additionally, the activating and inhibiting reactions of Figure 2C do not need to represent the same molecular mechanisms. For example, the generation of G6P is driven by the phosphorylation of glucose by HK2, whereas glucose induces HK2 generation by activating the HK2 promoter. We use the system in Figure 2C as a simplified representation of glucose uptake in cancer, and as a basis for our mathematical model.

**Figure 2:**
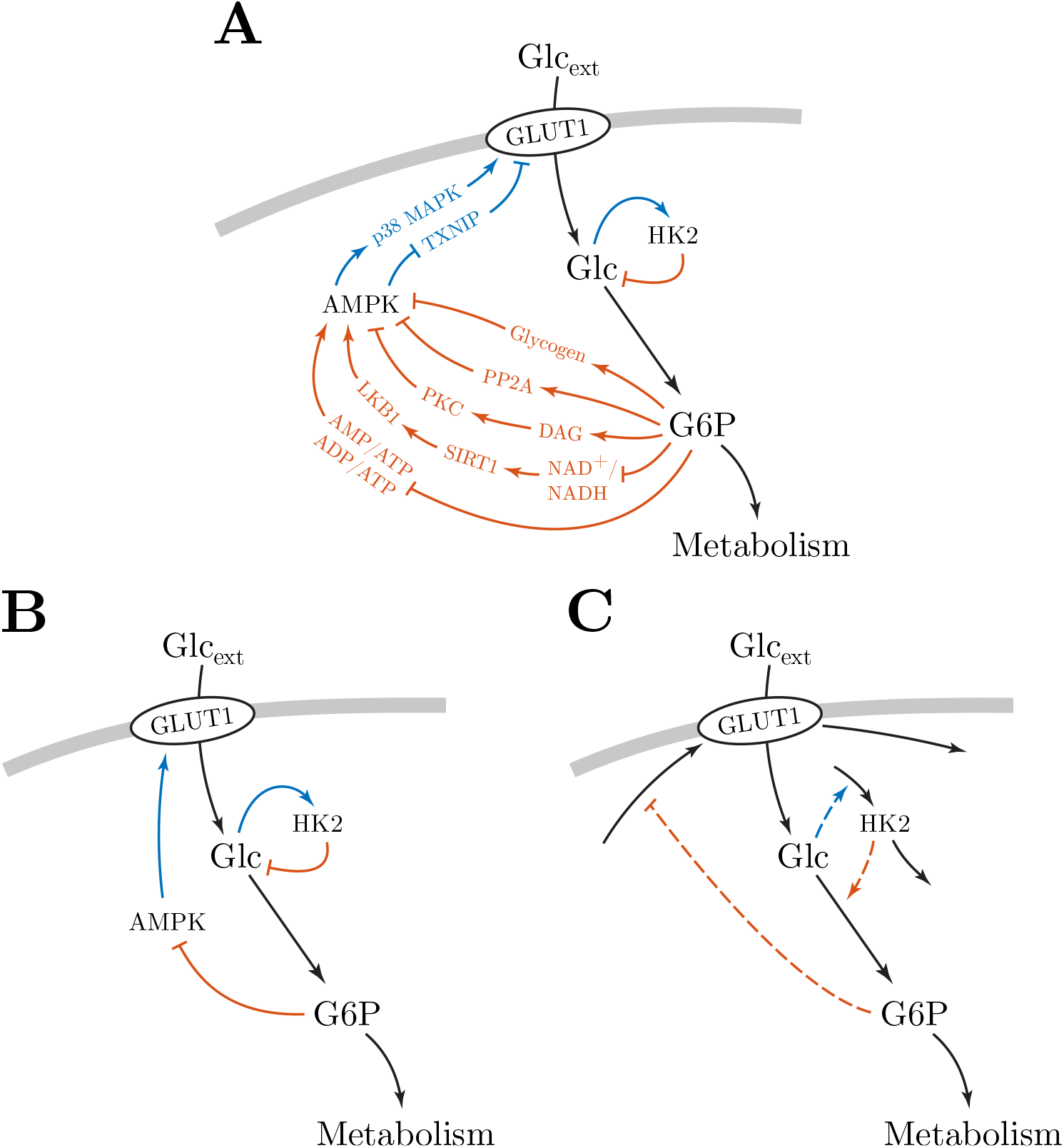
Panel A summarizes the control mechanisms of glucose uptake in cancer reported in the literature. Line marker-ends indicate the effect one compound has on another, arrowhead for positive and flat head for negative. Colored pathways indicate the overall effect of that pathway, blue for positive and red for negative. Black lines represent the flow of glucose to metabolism. Panel B shows colored pathways grouped together based on similar overall effects. Panel C shows the system in panel B translated into a form where activating and inhibiting effects act on reactions generating and turning over compounds. This allows for the system to be described by a simplified mathematical model using reaction kinetic equations.

## METHODS

### Differential Gene Expression

Expression Atlas was used to collect differential gene expression data comparing cancer cells with normal (i.e. non-cancerous) cells, across a variety of tissues and cell types. Expression Atlas is an open science resource providing information on gene and protein expression in animal and plant samples of different cell types, organism parts, developmental stages, diseases, and other conditions (66). Expression Atlas contains thousands of selected microarray and RNA-sequencing datasets that are manually curated, annotated, checked for high quality, and processed using standardized analysis methods (66). For genes of interest, users can view baseline expression in tissues, and differential expression for biologically meaningful pairwise comparisons (66).

Differential expression data of the *SLC2A* gene family, *HK1-3*, *GCK*, *PFKM*, *PFKP*, *PKM*, and *PKLR* genes in human was gathered from the Expression Atlas database. We curated the data to ensure only experiments comparing cancer cells with normal cells were included. Differential gene expression experiments with drug treatments were removed. Expression Atlas reports experiment results as log_2_-fold changes. In this paper, we report the arithmetic mean of log_2_-fold changes (i.e. log_2_ of the geometric mean fold change) for each gene across all experiments. Additional genes were analyzed, but due to low number of experiments (less than 5), are not included in the main results (see Section S1 in the Supporting Material).

### Computational Methods

Systems of ODEs are solved numerically in initial value problems using Matlab R2018a and the ode45 ODE solver, based on the Dormand-Prince (4, 5) pair (67). Initial values and parameters for all simulations are provided in the Supporting Material. Simulation results are given in arbitrary units (arb. unit). Reaction rates are expressed as concentrations per unit of time.

## RESULTS AND DISCUSSION

### Corroborating the Reported Rewiring of Glycolysis in Cancer

Average log_2_-fold changes for key genes associated with glucose uptake and glycolysis, across a variety of tissues and cell types, are shown in Figure 3. The differential gene expression data largely corroborates the reported rewiring of glycolysis in cancer discussed above. Namely, a shift towards GLUT1 (*SLC2A1* gene) mediated glucose uptake, predominant expression of the PKM2 (PKM1 and PKM2 are different splicing products of the *PKM* gene (21)) isoform, and overexpression of HK2. We also found a slight upregulation of the *PFKP* gene in cancer, consistent with previous studies (34). Hence, the model proposed in Figure 2C appears to include the key components of glucose uptake in cancer, and provides a good basis for mathematical modeling.

**Figure 3:**
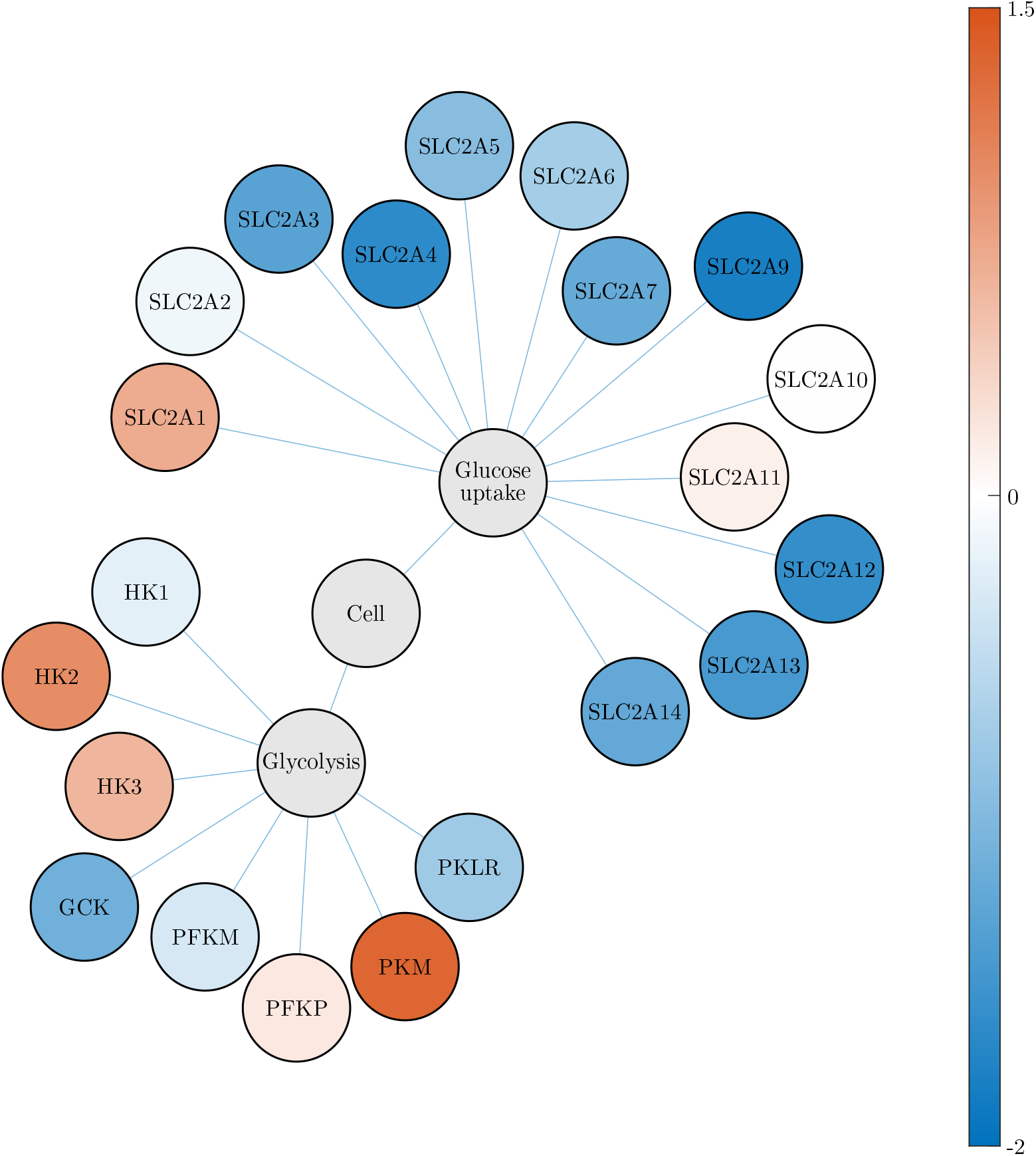
Differential gene expression of key genes associated with glucose uptake and glycolysis. The differential gene expression compares cancer cells with normal cells, across a variety of tissues and cell types, reported as the average log_2_-fold change of several experiments. Upregulation in cancer cells is indicated by red, and downregulation by blue. White indicates no change. The differential gene expression corroborates the reported rewiring of glycolysis in cancer. Namely, a shift towards GLUT1 (*SLC2A1* gene) mediated glucose uptake, overexpression of HK2, and a predominant reliance on PKM2 (PKM1 and PKM2 are different splicing products of the *PKM* gene (21)). We also found upregulation of the *HK3* and *PFKP* genes. See Section S1 in the Supporting Material for information on the individual differential gene expression experiments.

The results also shows an increased *HK3* transcript abundance in cancer. This is not surprising, since it has been shown that HK3 is upregulated by hypoxia, partially through HIF dependent signaling (68). Whereas HK2 bind to the outer mitochondrial membrane, HK3 does not (13, 68). A consequence of mitochondrial bound HK2 is the prevention of cell death by inhibiting formation of the mitochondrial permeability transition pore (MPTP) complex (13, 68). On the other hand, HK3 overexpression promotes cell survival in response to oxidative stress, decreases the production of reactive oxygen species (ROS), perserves mitochondrial membrane potential, and promotes mitochondrial biogenesis (68). Therefore, it is likely that HK2 and HK3 serve different, but complementary, roles in maintaining a highly glycolytic phenotype and promoting cancer cell survival. Notably, inhibition of glucose or G6P binding to the regulatory half of HK3 (N-terminal domain) impairs catalysis in the catalytic half (C-terminal domain), suggesting a cooperative effect of glucose binding in the regulatory half to subsequent binding in the catalytic half (68). Hence, it appears that HK3 interacts with glucose in a similar way to that of HK2 in Figure 2. As a result, we will only consider HK2 in the following mathematical modeling, but note that HK2 can be thought of as a pool of both HK2 and HK3.

### Modeling Rate Expressions in a Changing Volume

When modeling rate expressions in a changing volume, care must be taken so that concentrations are handled in the correct way. As an example we show how this is done for a simple enzyme reaction. The Michaelis–Menten equation describes the rate of an enzyme reaction, assuming steady state for the substrate-enzyme complex (69)

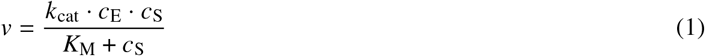

where *v* is the reaction rate, *k*_cat_ is the catalytic constant (or turnover number), *K*_M_ is the Michaelis constant, *c*_E_ is the (total) concentration of enzyme, and *c*_S_ is the concentration of substrate. We start by considering some compound x in a changing volume. Using the product rule, we express the change in concentration of x as

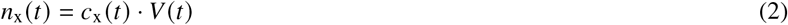

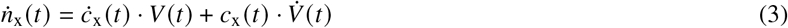

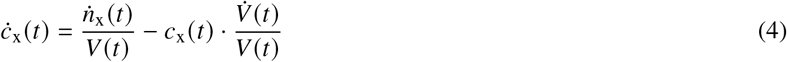

where *n*_x_ is the amount of compound, *c*_x_ is the concentration of compound, and *V* is the volume. We use dot notation to indicate time derivative. The first term of Eq. 4 is identical to the time derivative of *c*_x_ in a constant volume, while the second term represents the dilution of *c*_x_ (10). We will call this the *dilution term*. Using Eq. 1, we express the differential equation of a product P being formed by an enzyme reaction in a changing volume by introducing the dilution term from Eq. 4 (5)

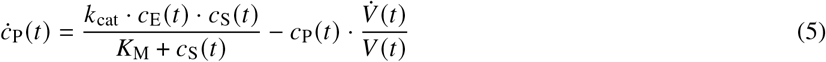

It is important to note that all constituents of the enzyme reaction, i.e. product, enzyme, and substrate, are diluted as the volume increases. This means that even if an enzyme is present in constant amount, a large enough volume increase can effectively stop the enzyme reaction by dilution of the enzyme concentration (5).

### Modeling Glucose Uptake in Cancer

We construct the mathematical model of glucose uptake in three steps, starting with glucose uptake and supply to metabolism without any of the control mechanisms discussed above. This system is shown in Figure 4A. Assuming low intracellular concentration of glucose due to rapid conversion by HK2, facilitated diffusion of glucose by GLUT1 can be approximated by the Michaelis–Menten equation (57, 58). In this first step of model construction, we assume that HK2 is not being generated and turned over (and therefore, HK2 synthesis is not activated by intracellular glucose), and that the concentration of HK2 simply dilutes as the cellular volume increases. The phosphorylation of glucose to G6P is modeled by the Michaelis–Menten equation, and the sink reaction to metabolism is modeled by a first order reaction with rate constant *k*_metabolism_. GLUT1 is assumed to be generated and turned over in reactions driven by enzymes E_1_ and E_2_, respectively, but feedback inhibition from G6P is omitted for the time being. The enzymes E_1_ and E_2_ are themselves present in constant amounts only (i.e. their concentrations simply dilute with increasing volume). We assume the production of GLUT1 is proportional to the concentration of E_1_, and that the degradation of GLUT1 by E_2_ is given by a Michaelis–Menten-type process. We are considering a growing cell, which introduces the dilution term from Eq. 4. The dynamical model is given by the following system of ODEs

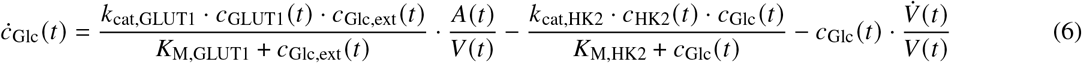

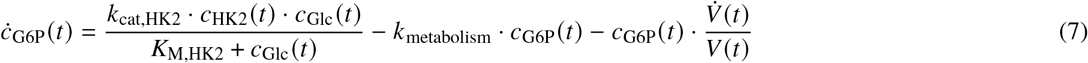

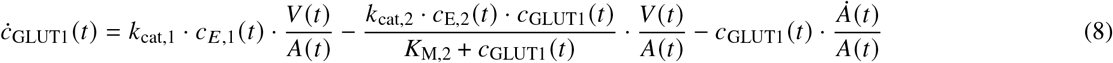

where *c*_Glc_ and *c*_G6P_ are concentrations in the cellular volume *V*, whereas *c*_GLUT1_ is a concentration at the cell surface *A*. As a consequence, the import of glucose is converted by the factor 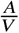 to a flux given with respect to the cellular volume. Similarly, the generation and degradation of GLUT1 are converted by the factor 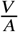 to fluxes with respect to the cell surface area, since the enzymes generating and turning over GLUT1 are situated inside the cell. HK2, E_1_, and E_2_ are not assumed to be generated and turned over, and their concentrations dilute from some initial concentrations as the cellular volume increases. These concentrations are given by *c*_x_ (*t*) = *n*_x_ /*V* (*t*) (x = HK2, E_1_, E_2_), where *n*_x_ is the amount of compound x (constant quantities).

**Figure 4:**
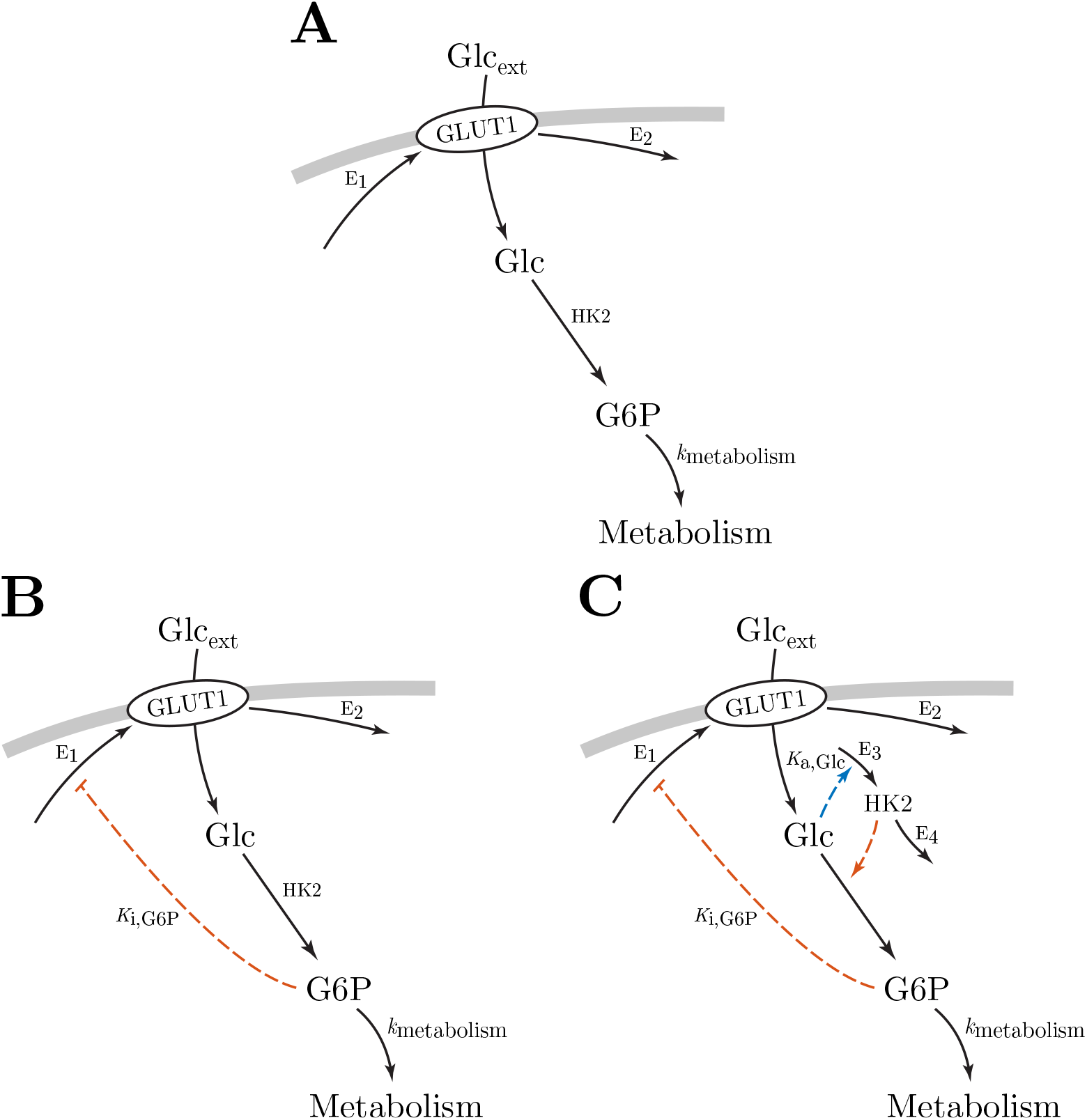
The mathematical model of glucose uptake is constructed in three steps. Panel A shows the first step, with only the uptake and supply of glucose to metabolism. The second step is shown in panel B, which includes feedback inhibition from G6P to GLUT1 mediated glucose uptake. Panel C shows the final step, where the model also includes the stabilizing feedback connection formed by HK2 and intracellular glucose.

We call the system of ODEs given by Eqs. 6–8 *model A*, corresponding to the system shown in Figure 4A. This model only describes the uptake of glucose and supply to metabolism, without any control mechanisms in place. To examine the regulatory mechanisms of glucose uptake, we build on this model, and add feedback inhibition from G6P to GLUT1 production. This feedback is based on the many pathways that regulate GLUT1 mediated glucose uptake via AMPK, summarized in Figure 2A. This way, a reduction in G6P level will reduce inhibition of GLUT1 production, thereby increasing GLUT1 mediated glucose uptake. We model this feedback by allosteric inhibition (specifically, a special case of mixed inhibition) of the reaction producing GLUT1 (69, 70). The model is shown in Figure 4B, and given by Eqs. 6–7, and the additional ODE

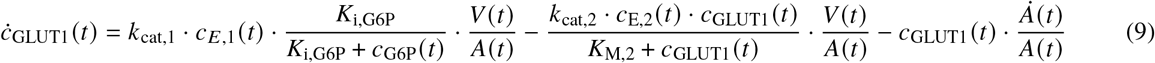

where *K*_i,G6P_ is the inhibition constant for the allosteric inhibition of GLUT1 production by G6P. We call this *model B*.

We simulate models A and B in three phases (Figure 5): In the first phase (white area, *t* = [0, 50]), the cellular volume is kept constant. In the second phase (light gray area, *t* = [50, 100]), we still maintain a constant cellular volume, and probe the regulatory function of the feedback inhibition in model B by reducing the extracellular glucose concentration by 75% at the start of the phase. In the third phase (dark gray area, *t* = [100, 150]), we investigate the effect of dilution on the two models by increasing the cellular volume linearly. The simulation results are shown in Figure 5, with initial values and parameters provided in Table S1 in the Supporting Material. Dashed red lines show the dynamical response of model A, and solid blue lines show model B. The bottom right plot of cellular volume (solid black line) and surface area (dashed black line) is the same for both simulations. In the first phase (white area), the cellular volume is constant and both systems have settled at steady state, producing a constant glycolytic flux (represented by the phosphorylation of glucose). At the start of the second phase (light gray area), extracellular glucose concentration is reduced while the cellular volume remains constant. Comparing the two models, we see that model A shows no adaptation to such a perturbation in glucose supply, resulting in reduced metabolite levels (intracellular glucose and G6P) and glycolytic flux. Model B, however, is able to fully compensate for the reduction in glucose supply. This is achieved by increasing the surface concentration of GLUT1, thereby increasing GLUT1 mediated glucose uptake to match the previous uptake rate of the system. In the final phase of the simulations (dark gray area), cellular volume starts to increase linearly. Although neither of the models are able to compensate for dilution, in the case of model B, GLUT1 production is increased in an attempt to mitigate the effect of dilution. However, this compensatory response is not being effectuated as the level of HK2 is not being maintained, resulting in reduced glycolytic flux as the concentration of HK2 dilutes. The simulation results show that feedback inhibition from downstream glycolytic metabolites to glucose transporters is sufficient for homeostatic control of glycolysis in a constant volume, consistent with principles of metabolic regulation by negative feedback (71, 72). The regulation of glucose transporters by feedback inhibition is, however, not sufficient to maintain a constant glycolytic flux during growth. It appears that control of HK2 may be necessary for homeostatic control during growth. In order to compensate for dilution, we extend model B, and add activation of HK2 synthesis by intracellular glucose. As mentioned earlier, this forms a stabilizing feedback connection together with the phosphorylation of glucose to G6P (62). This feedback connection stabilizes the level of HK2 in the presence of dilution, such that the concentration of HK2 remains more or less constant during growth, providing robustness to dilution in the first irreversible step of glycolysis. In turn, this robustness allows GLUT1 mediated glucose uptake to regulate the glycolytic flux during growth. With this addition it is necessary to add reactions generating and turning over HK2, so that activation by intracellular glucose can be mediated through these reactions. The activation of HK2 synthesis is modeled by allosteric activation (specifically, a special case of mixed activation) (69, 70).

**Figure 5:**
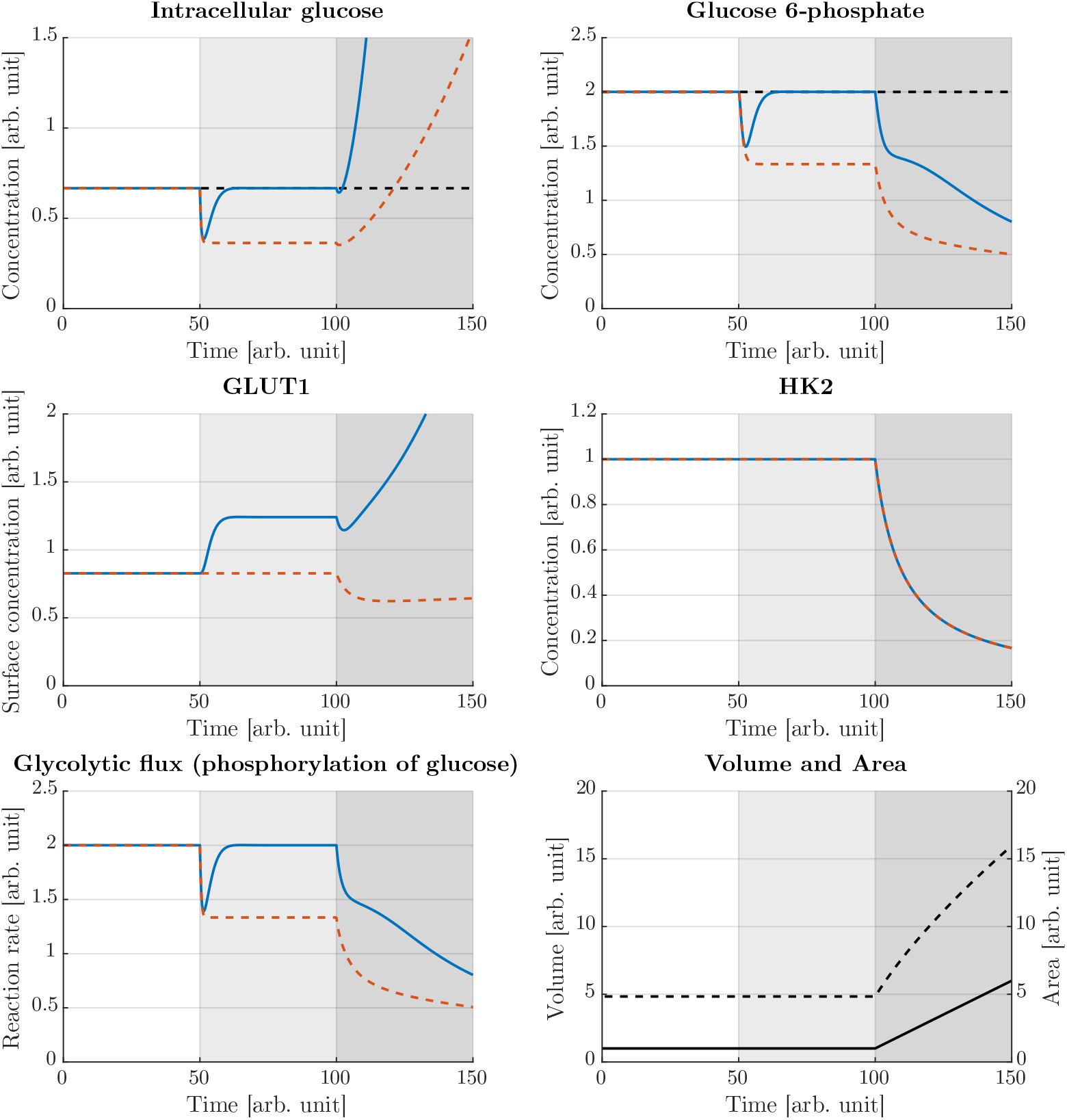
Simulation results of model A (dashed red lines) and model B (solid blue lines). The bottom right plot of volume (solid black line) and surface area (dashed black line) is the same for the two simulations. Initially, the cellular volume is kept constant and the systems have settled at steady state (white area, *t* = [0, 50]). At the start of the second phase (light gray area, *t* = [50, 100]), extracellular glucose concentration is reduced by 75%. Whereas model A shows no adaptation in this phase, model B is able to regulate intracellular glucose and G6P levels back to pre-perturbed values (dashed black lines), and maintain homeostatic control of glycolysis (regulation of the glycolytic flux). In the last phase (dark gray area, *t* = [100, 150]), the cellular volume starts to increase linearly. Neither of the models are able to compensate for dilution, however, GLUT1 mediated glucose uptake is increased in model B, but due to dilution of HK2 this compensatory response is not being effectuated. Note that due to dilution of HK2, intracellular glucose accumulates in the final phase. Such an accumulation can not go on forever, and as the concentration of intracellular glucose approaches that of extracellular glucose, our assumption of GLUT1 mediated glucose uptake following the Michaelis–Menten equation will break down. A similar limit for the surface concentration of GLUT1 will also likely be approached. Nevertheless, the simulation results are able to show that models A and B do not achieve homeostatic control of glysolysis in the presence of dilution. Initial values and parameters are provided in Table S1 in the Supporting Material.

We assume the synthesis and degradation of HK2 are driven by enzymes E_3_ and E_4_, respectively, where the synthesis is proportional to the level of E_3_, and the degradation by E_4_ follows a Michaelis–Menten-type process. The model is shown in Figure 4C and given by Eqs. 6–7, 9, and the following ODE describing the change in HK2 concentration

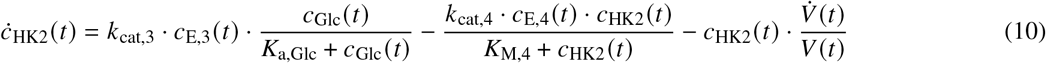

where *K*_a,Glc_ is the activation constant for the allosteric activation of HK2 synthesis by intracellular glucose. The enzymes E_i_ (i = 1,2,3,4) are not assumed to be generated and turned over, and their concentrations dilute as the volume increases. These concentrations are given by *c*_E,i_ (*t*) = *n*_E,i_ *V*(*t*) (i = 1, 2, 3, 4), where *n*_E,i_ (the amount of E_i_) are constant quantities. We call this *model C*.

We simulate model C in four phases (Figure 6): In the first phase (white area, *t* = [0,400]), the volume is kept constant. In the second phase (light gray area, *t* = [400,800]), we increase the volume linearly to investigate whether model C is able to maintain homeostatic control of glycolysis during growth. While the volume is still increasing, extracellular glucose concentration is increased 4-fold at the start of the third phase (dark gray area, *t* = [800,1200]). Finally, in the last phase (white area, *t* = [1200,1600]), volume increase is stopped. The simulation results are shown in Figure 6, with initial values and parameters provided in Table S2 in the Supporting Material. The bottom right plot shows volume (solid black line) and surface area (dashed black line) during the simulation. In the first phase (white area), the system has settled at steady state, producing a constant glycolytic flux. In the second phase (light gray area), we see that model C is able to compensate for dilution and produce a constant glycolytic flux, however, steady state values are shifted compared to steady state values without growth (dashed black lines in intracellular glucose and G6P plots). These growth associated offsets could indicate the inability of the control mechanisms to fully compensate for dilution. However, they could also represent set-point changes during the growth phase. To investigate the cause of the growth associated offsets, we increase extracellular glucose concentration as the cellular volume is growing. This is done at the start of the third phase (dark gray area). Due to the subsequent increase in glucose uptake, a sudden reduction in the surface concentration of GLUT1 follows. Nevertheless, the surface concentration of GLUT1 continues to increase throughout this phase in order to compensate for dilution. Interestingly, it seems that the control mechanisms attempt to bring the system back to steady state values associated with growth, not steady state values associated with constant volume. If the latter were true, we would not expect to see the regulatory action in Figure 6 bringing intracellular glucose and G6P levels away from steady state values associated with constant volume (dashed black lines). This suggests that the growth associated offsets may be caused by set-point changes, rather than the inability to maintain the glycolytic flux during growth. Finally, the first phase is repeated and the volume is kept constant again, but is now much larger (white area). In this phase, we see that metabolite levels and the glycolytic flux return to steady state values associated with constant volume. This is achieved by the increase of surface concentration of GLUT1 during the growth phase, which is made possible due to the relationship between cellular volume and cell surface area (see Section S2 in the Supporting Material). Thus, the growth associated offsets appears to be dependent on the rate of volume increase, not the total volume.

**Figure 6:**
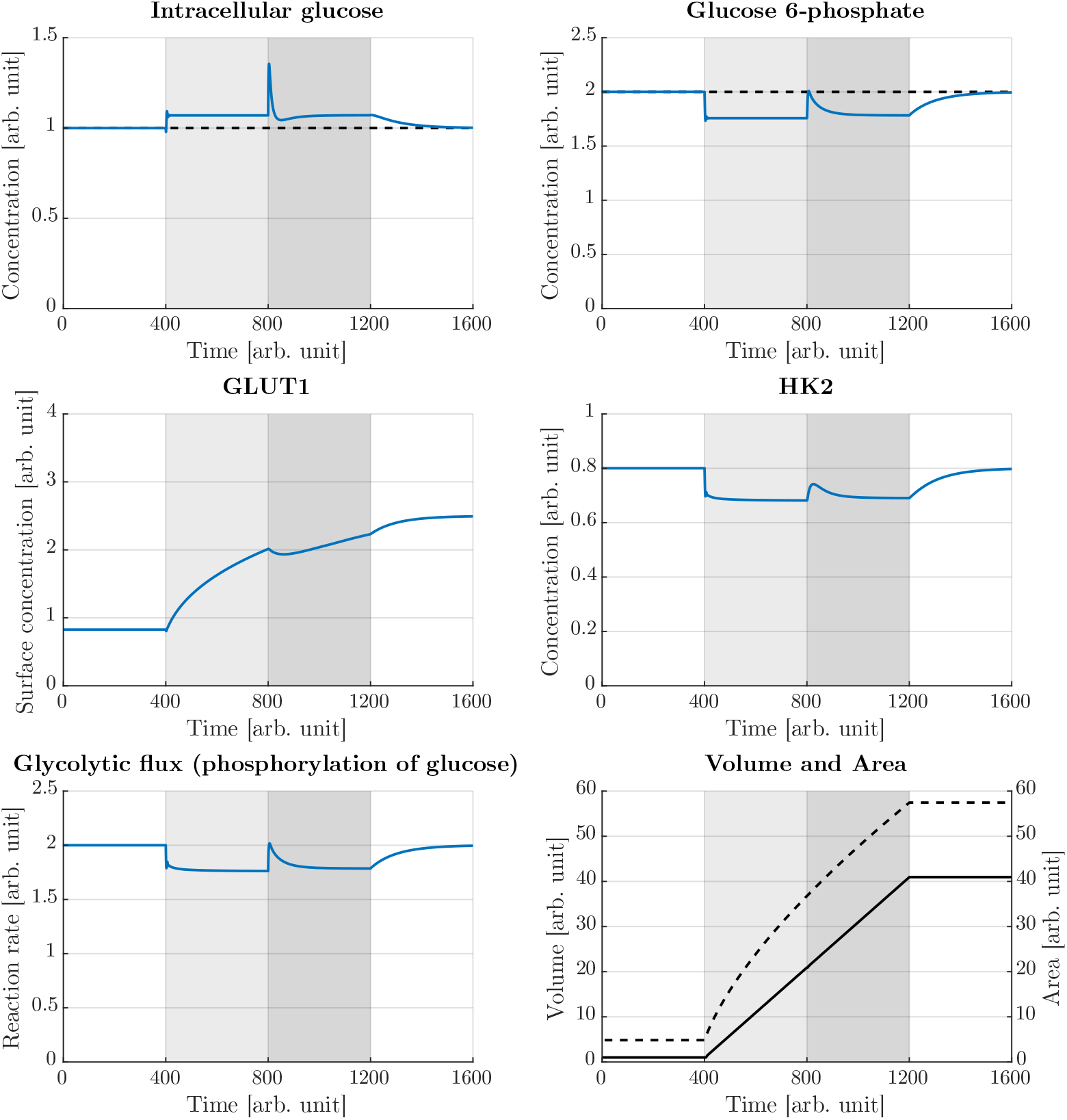
Simulation results of model C. The bottom right plot shows volume (solid black line) and surface area (dashed black line) during the simulation. Initially, the cellular volume is kept constant, and the system has settled at steady state (white area, *t* = [0, 400]). In the second phase (light gray area, *t* = [400, 800]), the cellular volume increases linearly. During this phase, metabolite levels and the glycolytic flux are maintained at constant levels, however, with offsets from steady state values associated with constant volume (dashed black lines in intracellular glucose and G6P plots). At the start of the third phase (dark gray area, *t* = [800, 1200]), as the volume is still increasing, the concentration of extracellular glucose is increased 4-fold. Interestingly, the system is regulated back to steady state values associated with growth, and away from steady state values associated with constant volume. This suggests that the growth associated offsets are caused by set-point changes. Finally, the first phase is repeated, and the volume is kept constant (white area, *t* = [1200, 1600]). In this phase we see that metabolite levels and the glycolytic flux return to steady state values associated with constant volume. This indicates that the growth associated offsets are dependent on the growth rate, rather than the total volume. Initial values and parameters are provided in Table S2 in the Supporting Material.

Taken together, the simulation results of models A, B, and C in Figure 5 and Figure 6 show that while negative feedback from downstream metabolites to nutrient transporters is sufficient for homeostatic control in a constant volume, it is necessary to stabilize the concentrations of intermediate enzymes in order to achieve homeostatic control during growth. The simulations also demonstrate that during the growth phase, growth associated offsets from steady state values associated with constant volume are observed, and that these offsets appears to be caused by set-point changes that are dependent on the growth rate. Importantly, investigations into control mechanisms similar to the ones identified in this paper, have shown that growth associated offsets become negligible if the kinetics of the controller species behave on a timescale much faster than cell growth (9, 10, 73–75). These control mechanisms, called ICMs, achieve robust homeostatic control due to a negative feedback structure that includes integral action. In the following, we take a closer look at the function of such ICMs and show how the control mechanisms discussed above realize integral action and dilution resistance.

### A Closer Look at Integral Control Motifs

Asymptotic regulation is the notion that a regulation error approaches zero, i.e. the output perfectly reaches a desired reference, as time tends to infinity (76). If asymptotic regulation is achieved in the presence of disturbances, asymptotic disturbance rejection (also called robustness) is achieved (76). In the case of constant reference signal, or set-point, and constant disturbance, asymptotic regulation and disturbance rejection can be achieved by integral action (76). A block diagram of negative feedback with integral action is shown in Figure 7. For a system subject to disturbance *w*, the output *y* is to be regulated to a set-point *r*. This is achieved by comparing the system output to the set-point, giving the regulation error *e* = *r − y*. The integral controller integrates the regulation error, producing the control action input *u* to the system. Thus, when the system output deviates from the set-point, the regulation error is non-zero, which produces a change in the control action. Because the feedback is negative, this change in control action counteracts the deviation of the system output from the set-point. Importantly, when the system output reaches the desired set-point, the output is maintained exactly at the set-point, as the “memory” element of the integral controller stores the accumulated regulation error (9). The block diagram in Figure 7 suggests a constant integral gain *G*_i_, though this gain can be variable, often referred to as gain scheduling (62, 76). The mathematical description of the integral controller, called the integral control law, is given by

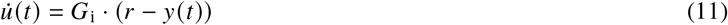

**Figure 7:**
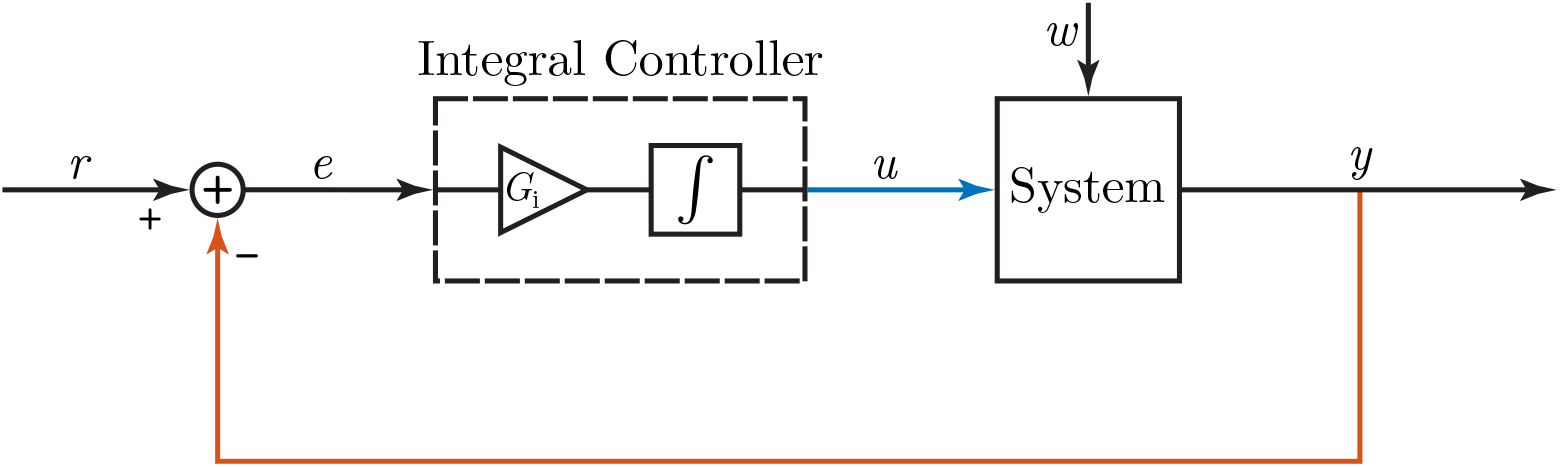
Block diagram of negative feedback with integral action. The system output *y* is fed back (red output feedback) and compared to the reference signal *r* to produce the regulation error *e* = *r − y*. The regulation error is multiplied by an integral gain *G*_*i*_ and integrated over time to produce the control action *u* (blue system input). In the presence of an uncontrolled disturbance *w* (black disturbance input), a deviation in system output from the reference will cause a non-zero regulation error. This produces a change in the control action, and since the feedback is negative, this control action functions to contract the deviation in system output from the reference.

Investigations into robust homeostatic systems have revealed several motifs that include negative feedback with integral action (9, 62, 77, 78). The control mechanisms considered in this paper correspond to a class of ICMs called homeostatic controller motifs (62). It has been shown that these homeostatic controller motifs are robust to all parameter perturbations that do not destroy the stability of the system (79). For example, feedback inhibition from G6P to GLUT1 generation produces the same structure as that of Figure 7. Red output feedback corresponds to the inhibition of GLUT1 generation by G6P, the integral controller block corresponds to GLUT1 level, blue system input corresponds to GLUT1 mediated glucose uptake, and the system block corresponds to the level of G6P. The black disturbance input corresponds to perturbations made in extracellular glucose and cellular volume. By manipulating Eq. 9, we show that GLUT1 functions as an integral controller for G6P level

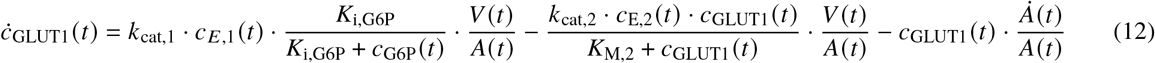

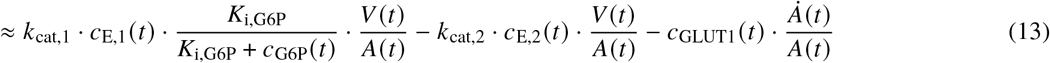

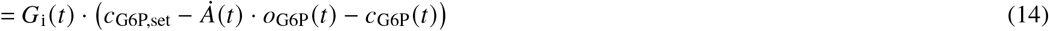

where we make the simplification *K*_M,2_ ≪ *c*_GLUT1_. The following definitions are made

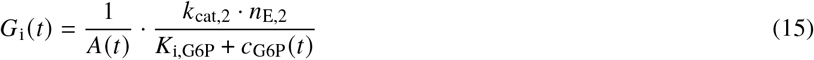

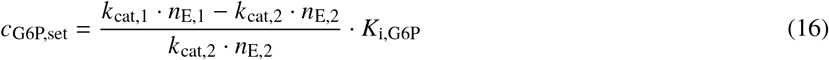

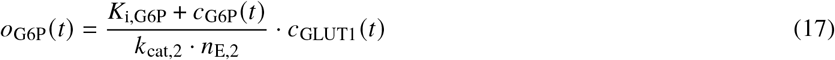

The set-point for G6P level, *c*_G6P,set_, is given entirely by parameters associated with GLUT1 generation and degradation. This means that perturbations in G6P are fully compensated for, as the set-point remains unchanged for such perturbations (73). In the case without growth, *Ȧ* = 0, Eq. 14 is reduced to the same form as the integral control law in Eq. 11. In the case with growth, a growth associated offset, *Ȧ* ‧ *o*_G6P_, is introduced. However, if the reaction rates for the generation and degradation of GLUT1 behave on a timescale much faster than the rate of dilution, *o*_G6P_ is small, and the growth associated offset becomes negligible. ICMs with controller reactions much faster than the rate of dilution are called quasi-ICMs, and characteristically show small growth associated offsets (9). One mechanism of regulating GLUT1 and HK2 activity is through translocation between biological membranes and the cytosol, indicating that the activity of these species can respond quickly, and that regulation of glucose uptake in cancer may achieve dilution resistance through the formation of quasi-ICMs (39, 55, 80). Similar to GLUT1, it is possible to show that HK2 functions as an integral controller for the level of intracellular glucose (See Section S3 in the Supporting Material).

## CONCLUSION

In this paper, we have constructed a mathematical model of glucose uptake based on the reported rewiring of glycolysis in cancer and differential gene expression of cancer and normal cells. With basis in the literature, we added control mechanisms to the model in a stepwise manner, in order to investigate the role each regulatory mechanism serve. Expectedly, we found that feedback inhibition from downstream glycolytic metabolites to glucose transporters is sufficient for homeostatic control of glycolysis in a constant volume. However, in a growing volume, we found that regulation of intermediate glycolytic enzymes is needed for homeostatic control of glycolysis. Cancer cells show a shift towards GLUT1 mediated glucose uptake and a reliance on HK2. We found that these species form regulatory mechanisms for glycolysis through their interactions with glycolytic metabolites. These regulatory mechanisms are a class of ICMs known as homeostatic controller motifs, and achieve robust homeostatic control by negative feedback with integral action (9, 62). Our simulation results show that during growth, offsets from steady state values associated with constant volume are observed. These growth associated offsets can be interpreted as set-point changes, and are dependent on the growth rate of the cell, not the total volume. In his definition of homeostasis, Cannon emphasized that homeostasis does not imply perfect adaptation to disturbances, but allows for some variability in steady state (81). Similarly, rheostasis is defined as systems that show homeostatic control at any one instant, but over the span of time show change in the regulated level (82). Living organisms are not necessarily concerned with perfect regulation, but rather with the presence of some level of regulation. Thus, it is likely that sufficient regulation can be achieved even if the growth associated offsets are fairly large. This variability in steady state can then be viewed as a relaxing condition on the control mechanisms employed (83).

Investigations into ICMs have shown that growth associated offsets becomes negligible if the rates of the controller reactions, the generation and degradation of GLUT1 and HK2, are much faster compared to the rate of dilution (9, 10, 73–75). In our model, enzymes responsible for generating and removing the controller species (E_i_, i = 1,2,3,4) are present in constant amounts only, meaning that their concentrations simply dilute with increasing volume. This is a worst-case scenario in which regulation in the presence of dilution is possible. In this scenario, we found that model C is able to compensate for dilution in a linearly increasing volume. However, most protein and mRNA concentrations are independent of cell size, and therefore it is likely that the concentrations *c*_E,i_ (i = 1,2,3,4) should be considered constant (7). In this case, it has been shown that regulation is possible even in the presence of dilution in an exponentially increasing cell volume (9, 10).

Taking a closer look at feedback inhibition from downstream glycolytic metabolites to GLUT1 mediated glucose uptake, we have shown how this control mechanism realizes integral action to regulate glycolysis. We have also shown how dilution affects this ICM, and related the growth associated offset in G6P to a term dependent on the growth rate of the cell. In recent years, ICMs have garnered much attention (9, 10, 62, 75, 77, 78). This paper uses glucose uptake in cancer to demonstrate a systems biology approach to the analysis of complex regulatory systems, where control mechanisms are reduced into their essential components, which can then be represented by ICMs. A benefit of this approach is simplifying the mathematical description of the complex system, while retaining the essential behavior, such that a more manageable system can be considered and an in-depth analysis of its function can be done.

## Supporting information

Differential gene expression data

## AUTHOR CONTRIBUTIONS

DT, TD, KT, FF, and PR conceived and designed the research. DT and TD performed simulations. DT, TD, KT, and FF analyzed the data. DT and GF wrote the manuscript with contributions from all coauthors.

