## Supporting Material for "Exploring Mechanisms of Glucose Uptake Regulation and Dilution Resistance in Growing Cancer Cells"

### S1 Differential Gene Expression

Differential gene expression data collected from the Expression Atlas[1] database are provided in the file `Differential expression.xlsx`. The first sheet shows mean $\log_2$-fold change for the genes considered, along with the number of experiments used to calculate the mean. The following sheets show $\log_2$-fold change for each individual experiment, indicating the type of cells compared.

### S2 Note on Cellular Volume and Surface Area

Assuming spherical cells, the surface area $A$ and its rate of change $\dot{A}$ are calculated from the cellular volume $V$ and change in cellular volume $\dot{V}$ as follows

$$A(t) = 4\pi \left( \frac{3V(t)}{4\pi} \right)^{\frac{2}{3}} \tag{1}$$

$$\dot{A}(t) = \frac{2\dot{V}(t)}{\left( \frac{3V(t)}{4\pi} \right)^{\frac{1}{3}}} \tag{2}$$

This gives the relationship

$$\frac{\dot{V}(t)}{V(t)} = \frac{3}{2} \cdot \frac{\dot{A}(t)}{A(t)} \tag{3}$$

which expresses the difference in dilution between concentrations in the cellular volume and at the cell surface. This is the reason the surface concentration of GLUT1 does not settle at a constant level during growth (see Figure 6 in the main paper). In Eq. 14 in the main paper, the offset $\dot{A} \cdot o_{\mathrm{G6P}}$ does not fully account for the difference in G6P set-point and observed level during growth. In fact, the offset is off by a factor $\frac{3}{2}$, due to this difference in dilution of components. The same is not true for the negative feedback connection formed by HK2 and intracellular glucose, as both components are concentrations with respect to the cellular volume.

[1]`https://www.ebi.ac.uk/gxa/home` (see also reference (66) in the main paper)

### S3 Integral Control of Intracellular Glucose

Similar to how GLUT1 functions as an integral controller for G6P, HK2 functions as an integral controller for intracellular glucose

$$\dot{c}_{\mathrm{HK2}}(t) = k_{\mathrm{cat},3} \cdot c_{\mathrm{E},3}(t) \cdot \frac{c_{\mathrm{Glc}}(t)}{K_{\mathrm{a,Glc}} + c_{\mathrm{Glc}}(t)} - \frac{k_{\mathrm{cat},4} \cdot c_{\mathrm{E},4}(t) \cdot c_{\mathrm{HK2}}(t)}{K_{\mathrm{M},4} + c_{\mathrm{HK2}}(t)} - c_{\mathrm{HK2}}(t) \cdot \frac{\dot{V}(t)}{V(t)} \quad (4)$$

$$\approx k_{\mathrm{cat},3} \cdot c_{\mathrm{E},3}(t) \cdot \frac{c_{\mathrm{Glc}}(t)}{K_{\mathrm{a,Glc}} + c_{\mathrm{Glc}}(t)} - k_{\mathrm{cat},4} \cdot c_{\mathrm{E},4}(t) - c_{\mathrm{HK2}}(t) \cdot \frac{\dot{V}(t)}{V(t)} \quad (5)$$

$$= G_{\mathrm{i}}(t) \cdot \Big(c_{\mathrm{Glc,set}} - \dot{V}(t) \cdot o_{\mathrm{Glc}}(t) - c_{\mathrm{Glc}}(t)\Big) \quad (6)$$

where the simplification $K_{\mathrm{M},4} \ll c_{\mathrm{HK2}}$ is made. The following definitions are made

$$G_{\mathrm{i}}(t) = \frac{1}{V(t)} \cdot \frac{k_{\mathrm{cat},4} \cdot n_{\mathrm{E},4} - k_{\mathrm{cat},3} \cdot n_{\mathrm{E},3}}{K_{\mathrm{a,Glc}} + c_{\mathrm{Glc}}(t)} \quad (7)$$

$$c_{\mathrm{Glc,set}} = \frac{k_{\mathrm{cat},4} \cdot n_{\mathrm{E},4}}{k_{\mathrm{cat},3} \cdot n_{\mathrm{E},3} - k_{\mathrm{cat},4} \cdot n_{\mathrm{E},4}} \cdot K_{\mathrm{a,Glc}} \quad (8)$$

$$o_{\mathrm{Glc}}(t) = \frac{K_{\mathrm{a,Glc}} + c_{\mathrm{Glc}}(t)}{k_{\mathrm{cat},4} \cdot n_{\mathrm{E},4} - k_{\mathrm{cat},3} \cdot n_{\mathrm{E},3}} \cdot c_{\mathrm{HK2}(t)} \quad (9)$$

### S4 Initial Values and Parameters

| | **Initial Values** | |
|---|---|---|
| Name | Model A | Model B |
| $c_{\mathrm{Glc}}$ | 0.6669 | 0.6669 |
| $c_{\mathrm{G6P}}$ | 2.0004 | 2.0004 |
| $c_{\mathrm{GLUT1}}$ | 0.8273 | 0.8273 |
| | **Parameters** | |
| $n_{\mathrm{HK2}}$ | 1.0000 | 1.0000 |
| $V$ | $f(t) = 1.0000 + \dot{V} \cdot t$ | $f(t) = 1.0000 + \dot{V} \cdot t$ |
| $\dot{V}$ | $f(t) = \begin{cases} 0, & t < 100 \\ 0.1000 & t \geq 100 \end{cases}$ | $f(t) = \begin{cases} 0, & t < 100 \\ 0.1000 & t \geq 100 \end{cases}$ |
| $c_{\mathrm{Glc,ext}}$ | $f(t) = \begin{cases} 5.0000, & t < 50 \\ 1.2500, & t \geq 50 \end{cases}$ | $f(t) = \begin{cases} 5.0000, & t < 50 \\ 1.2500, & t \geq 50 \end{cases}$ |
| $k_{\mathrm{cat,GLUT1}}$ | 0.6000 | 0.6000 |
| $K_{\mathrm{M,GLUT1}}$ | 1.0000 | 1.0000 |
| $k_{\mathrm{cat,HK2}}$ | 5.0000 | 5.0000 |
| $K_{\mathrm{M,HK2}}$ | 1.0000 | 1.0000 |
| $k_{\mathrm{metabolism}}$ | 1.0000 | 1.0000 |
| $k_{\mathrm{cat},1}$ | 1.0000 | 6.0000 |
| $n_{\mathrm{E},1}$ | 1.0000 | 1.0000 |
| $k_{\mathrm{cat},2}$ | 2.0000 | 2.0000 |
| $K_{\mathrm{M},2}$ | 0.8273 | 0.0001 |
| $n_{\mathrm{E},2}$ | 1.0000 | 1.0000 |
| $K_{\mathrm{i,G6P}}$ | – | 1.0000 |

Table S1: Initial values and parameters for the simulations of model A and B in Figure 5.

| **Initial Values** | |
|---|---|
| Name | Model C |
| $c_{\text{Glc}}$ | 0.9997 |
| $c_{\text{G6P}}$ | 2.0004 |
| $c_{\text{GLUT1}}$ | 0.8273 |
| $c_{\text{HK2}}$ | 0.8002 |
| **Parameters** | |
| $V$ | $f(t) = 1.0000 + \dot{V} \cdot t$ |
| $\dot{V}$ | $f(t) = \begin{cases} 0.0500, & 400 \leq t < 1200 \\ 0, & \text{otherwise} \end{cases}$ |
| $c_{\text{Glc,ext}}$ | $f(t) = \begin{cases} 5.0000, & t < 800 \\ 20.0000, & t \geq 800 \end{cases}$ |
| $k_{\text{cat,GLUT1}}$ | 0.6000 |
| $K_{\text{M,GLUT1}}$ | 1.0000 |
| $k_{\text{cat,HK2}}$ | 5.0000 |
| $K_{\text{M,HK2}}$ | 1.0000 |
| $k_{\text{metabolism}}$ | 1.0000 |
| $k_{\text{cat,1}}$ | 6.0000 |
| $n_{\text{E,1}}$ | 1.0000 |
| $k_{\text{cat,2}}$ | 2.0000 |
| $K_{\text{M,2}}$ | 0.0001 |
| $n_{\text{E,2}}$ | 1.0000 |
| $k_{\text{cat,3}}$ | 2.0000 |
| $n_{\text{E,3}}$ | 1.0000 |
| $k_{\text{cat,4}}$ | 1.0000 |
| $K_{\text{M,4}}$ | 0.0001 |
| $n_{\text{E,4}}$ | 1.0000 |
| $K_{\text{i,G6P}}$ | 1.0000 |
| $K_{\text{a,Glc}}$ | 1.0000 |

Table S2: Initial values and parameters for the simulation of model C in Figure 6.
